# Characterisation of deubiquitylating enzymes in the cellular response to high-LET ionising radiation and complex DNA damage

**DOI:** 10.1101/453118

**Authors:** Rachel J. Carter, Catherine M. Nickson, James M. Thompson, Andrzej Kacperek, Mark A. Hill, Jason L. Parsons

## Abstract

**Purpose:** Ionising radiation, particular high linear energy transfer (LET) radiation, can induce complex DNA damage (CDD) where two or more DNA lesions are induced in close proximity which contributes significantly to the cell killing effects. However knowledge of the enzymes and mechanisms involved in co-ordinating the recognition and processing of CDD in cellular DNA are currently lacking.

**Methods and Materials:** An siRNA screen of deubiquitylation enzymes was conducted in HeLa cells irradiated with high-LET -particles or protons, versus low-LET protons and x-rays, and cell survival monitored by clonogenic assays. Candidates whose depletion led to decreased cell survival specifically in response to high-LET radiation were validated in both HeLa and oropharyngeal squamous cell carcinoma (UMSCC74A) cells, and the association with CDD repair was confirmed by using an enzyme modified neutral comet assay.

**Results:** Depletion of USP6 decreased cell survival specifically following high-LET α-particles and protons, but not by low-LET protons or x-rays. USP6 depletion caused cell cycle arrest and a deficiency in CDD repair mediated through instability of poly(ADP-ribose) polymerase-1 (PARP-1). This phenotype was mimicked using the PARP inhibitor olaparib.

**Conclusion:** USP6 controls cell survival in response to high-LET radiation by stabilising PARP-1 protein levels which is essential for CDD repair. We also describe synergy between CDD induced by high-LET protons and PARP inhibition in effective cancer cell killing.

## Introduction

DNA is the critical cellular target for ionising radiation (IR) and the induction of DNA double strand breaks (DSBs) but also complex (clustered) DNA damage (CDD) are thought to be the critical lesions contributing to the cell killing effects of IR. CDD is recognised as two or more DNA lesions induced in close proximity (e.g. within 1-2 helical turns of the DNA) and has been demonstrated to persist in cells and tissues several hours post-IR due to the difficulty in their repair (1,2). CDD formation increases with increasing linear energy transfer (LET) and been predicted by mathematical modelling to be important with proton beam irradiation, particularly at or around the Bragg peak where both high energy (or low-LET) protons and low energy protons (with increased LET) are generated (3-5). This has been shown indirectly by demonstrating that protons with increasing LET cause reduced cell survival (6,7) and increases in persistent DNA DSBs as revealed by 53BP1 foci (8). However, recent data from our laboratory has directly demonstrated using an enzyme modified neutral comet assay that low energy (relatively high-LET) protons generate significantly increased amounts of CDD compared to high energy (low-LET) protons or x-rays which persists for several hours post-irradiation (9).

Given that CDD is known to be important in the cell killing effects of IR, surprisingly the molecular and cellular mechanisms that respond to CDD within cellular DNA have been understudied. However we recently demonstrated that CDD induced by high-LET protons and α-particles causes elevations in the levels of histone H2B ubiquitylation on lysine 120 (H2B_ub_). We discovered that this is co-ordinated by the E3 ubiquitin ligases RNF20/40 and MSL2 that play important roles in the repair of CDD and in cell survival following high-LET protons. We postulated that this is a mechanism for enhancing CDD repair by promoting chromatin remodelling and/or actively recruiting DNA repair enzymes (9). Nevertheless this study has identified that ubiquitylation, particularly of histones, plays an important role in the cellular response to IR-induced CDD. Other DNA repair pathways, particularly DSB repair, are also known to be actively controlled by histone ubiquitylation that enhance DNA damage accessibility (10).

In addition to regulation of DNA repair via controlling chromatin accessibility, there are numerous studies demonstrating that DNA repair proteins themselves are subject to regulation by ubiquitylation, including those involved in DSB repair and in the repair of DNA base damage through the base excision repair pathway (10-12). This can be achieved by controlling DNA repair protein levels in response to the changing DNA damage environment and involves careful synchronisation of E3 ubiquitin ligases and deubiquitylation enzymes (DUBs) that control polyubiquitylation-dependent proteasomal degradation of the proteins. Given the essential role of ubiquitylation in co-ordinating the cellular DNA damage response, it can be hypothesized that DUBs will also play a central role following CDD induction by IR. However the specific DUBs that are responsive to high-LET irradiation that generates CDD in higher proportions compared to low-LET IR, have not been isolated to date. To this effect, we have utilised an siRNA screen targeting individual DUBs and analysed cell survival in response to low-versus high-LET IR. This approach has subsequently identified ubiquitin specific protease 6 (USP6) that modulates cell survival following high-LET IR by promoting efficient repair of CDD sites. We discovered that this is mediated through stabilisation of poly(ADP-ribose) polymerase-1 (PARP-1) and via cell cycle progression.

## Methods and Materials

### Antibodies and siRNA

The DUB siRNA library (ON-TARGETplus) containing pools of 4 individual siRNAs, and an individual siRNA targeting USP6 (USP6_13 5′-CAGCUAAGAUCUCAAGUCA-3′) were from Horizon Discovery (Cambridge, UK). The non-targeting control siRNA (AllStars Negative Control siRNA) was from Qiagen (Manchester, UK). The following antibodies were used:-USP6 and PARP-1 (both Santa Cruz Biotechnology, Heidelberg, Germany), phospho(T68)-Chk2 (New England Biolabs, Hitchin, UK), XRCC1 and APE1 (kindly provided by G. Dianov), γH2AX (Merck-Millipore, Watford, UK), 53BP1 (Bethyl Labs, Montgomery, USA), and actin (Sigma-Aldrich, Gillingham, UK).

### Cell culture

HeLa and UMSCC74A cells were cultured under standard conditions as previously described (13). siRNA knockdowns were performed for 48 h using Lipofectamine RNAiMAX (Life Technologies, Paisley, UK).

### Irradiation sources

Irradiation sources are as previously described (9). Briefly, cells grown in 35 mm dishes were exposed to low-LET x-rays (100 kV) using a CellRad x-ray irradiation (Faxitron Bioptics, Tucson, USA). Proton irradiations were performed at the Clatterbridge Cancer Centre and cells were irradiated directly by a ~1 keV/µm pristine beam of 58 MeV effective energy. Alternatively cells were irradiated using a modulator to generate a 27 mm spread-out Bragg peak (SOBP) and a 24.4 mm Perspex absorber to position the cells at the distal edge of the SOBP, corresponding to a mean proton energy of 11 MeV at a dose averaged LET of 12 keV/µm. Cells were irradiated with 3.26 MeV α-particles (LET of 121 keV/µm; dose rate of ~1.2 Gy/min) using a ^238^Pu irradiator. Cell cycle analysis was performed by fluorescence-activated cell sorting, as also previously described (9).

### Western blotting.

Whole cell extracts were prepared, separated by SDS-PAGE and analysed by quantitative Western blotting using the Odyssey image analysis system (Li-cor Biosciences, Cambridge, UK), as previously described (13,14).

### Clonogenic assays

Following irradiation in 35 mm dishes, cells were trypsinised, counted and a defined number seeded in triplicate into 6-well plates (250/500 for HeLa and 2000/4000 for UMSCC74A for unirradiated controls). Plating efficiencies for HeLa and UMSCC74A were ~40 % and 5 %, respectively and increasing cell numbers were used for increasing IR to account for these. Colonies were allowed to grow (~7-10 days), fixed and stained with 6 % glutaraldehyde, 0.5 % crystal violet for 30 min, and colonies counted using the GelCount colony analyser (Oxford Optronics, Oxford, UK). Surviving fraction (SF) was expressed as colonies per treatment versus colonies in the untreated control from at least three independent experiments, apart from the DUB siRNA screen which was from a single experiment.

### Enzyme modified neutral comet assay.

Detection of CDD using the enzyme modified neutral comet assay was as previously described (9). % Tail DNA values were calculated from at least three independent experiments.

### Immunofluorescent staining

Measurement of DNA repair protein foci (γH2AX and 53BP1) were examined as previously described (13) and mean foci/cell calculated from at least three independent experiments.

## Results

### Screening for specific DUBs involved in response to high-LET IR

We utilised an siRNA screen for up to 84 DUBs (pools of four siRNA duplexes) and analysed cell survival following IR, with a focus on high-LET IR which causes CDD. Using high-LET α-particle irradiation, depletion of only four DUBs (USP6, USP21, USP36 and DUB3) further reduced cell survival (by >50 %) compared to mock transfected cells (**Fig. 1A, E1A** and **Table E1**), which was a different subset of DUBs whose depletion reduced cell survival following low-LET x-ray irradiation (**Fig. 1B, E1A** and **Table E1**). Using high-LET protons (**Fig. 1C, E1B**) depletion of USP6 (plus CYLD and USP7) also reduced cell survival by >40 %. Interestingly a large number of DUBs (14 in total) caused enhanced radiosensitivity (>50 %) following low-LET protons. A complementary overexpression screen (**Fig. E2A** and **E2B**) revealed that overexpression of USP6 caused significantly increased cell resistance to α-particle irradiation, but had no impact on x-ray irradiation.

**Fig. 1.**
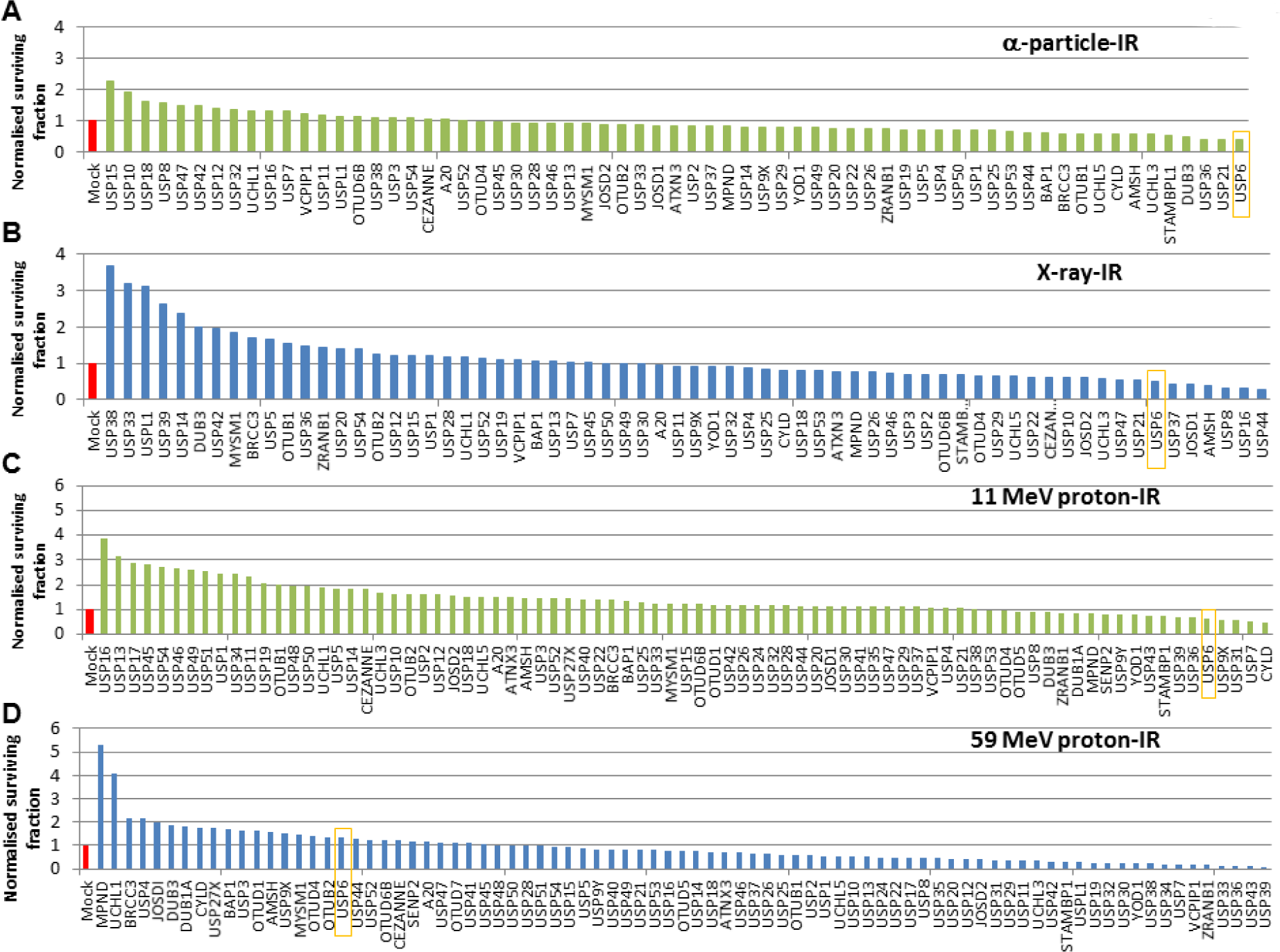
Screening of DUBs involved in cell survival following high‐ and low-LET irradiation. HeLa cells were treated with a pool of four siRNA oligonucleotides targeting individual DUBs for 48 h, and irradiated with (A) 0.5 Gy α-particles, (B) 1 Gy x-rays, (C) 2 Gy high-LET protons or (D) 2 Gy low-LET protons. Clonogenic survival of cells was analysed from a single experiment (using triplicate samples) and normalised against the mock treated control (red bar) which was set to 1.0 (equivalent to ~40 % cell survival post-irradiation).

Screening results were validated demonstrating that depletion of USP6 using a dose titration of low-LET x-rays (**Fig. 2A, 2C**) or low-LET protons (**Fig. 2E, 2G**) had no impact on radiosensitivity of HeLa cells in comparison to non-targeting (NT) control siRNA treated cells. In contrast, absence of USP6 caused a significant decrease in cell survival versus NT control siRNA cells following high-LET α-particle irradiation (**Fig. 2B, 2D**) and high-LET protons (**Fig. 2F, 2H**). Experiments were reproduced using a single siRNA sequence effective at supressing USP6 protein levels (**Fig. 3A**) that confirmed no effect on cellular radiosensitivity in response to low-LET protons (**Fig. 3B, 3D**) but which significantly reduced cell survival in response to high-LET protons compared against NT control siRNA cells (**Fig. 3C, 3E**). Depletion of USP6 was also able to specifically radiosensitise head and neck squamous cell carcinoma cells (UMSCC74A) to high-LET protons, but not low-LET protons (**Fig. E3A-D**).

**Fig. 2.**
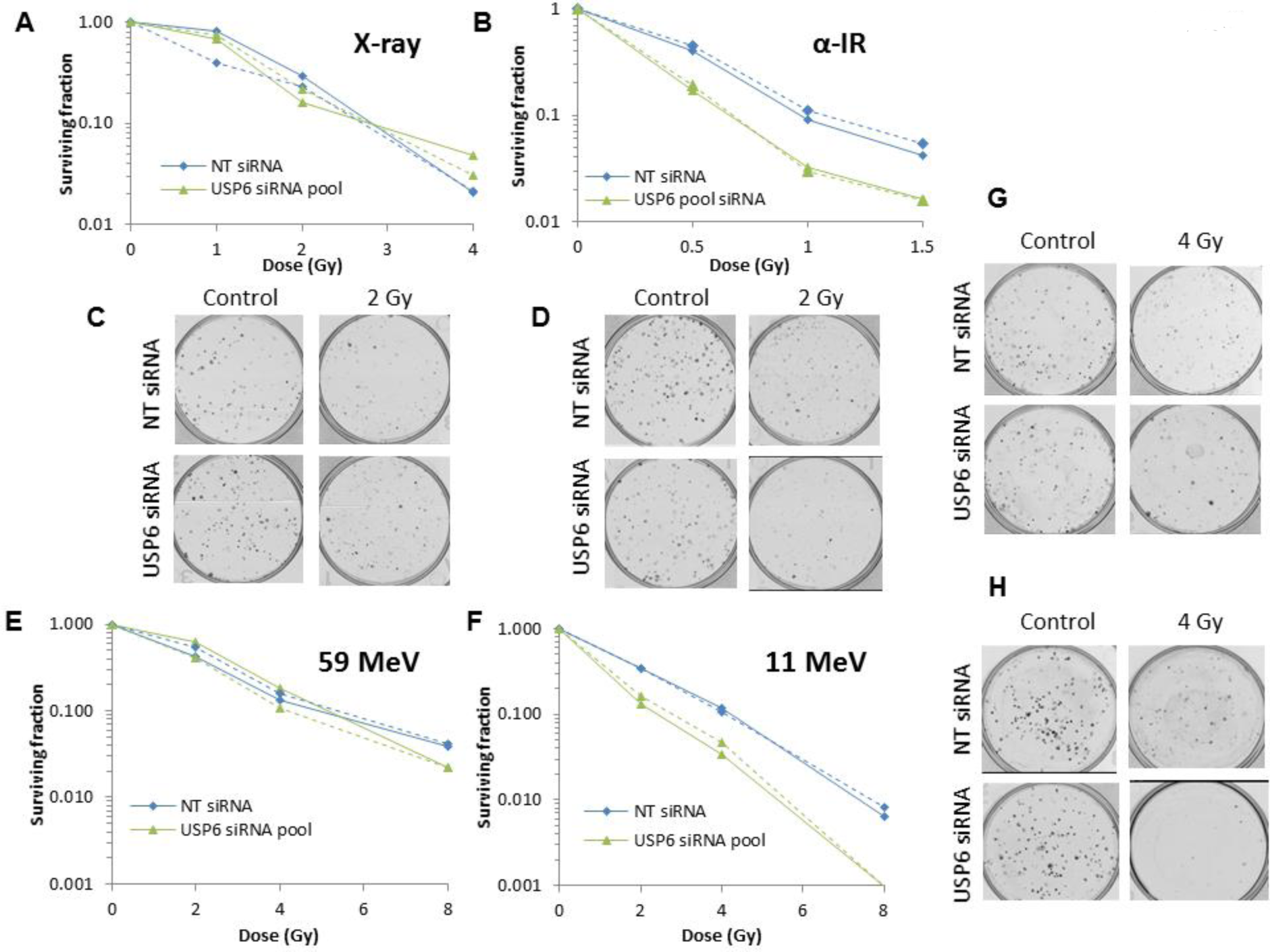
Validation of USP6 in controlling radiosensitivity in response to high-LET α-particles and protons. HeLa cells were treated with a pool of four siRNAs targeting USP6, or a non-targeting (NT) control siRNA for 48 h, and irradiated with increasing doses of (A, C) x-rays, (B, D) α-particles, (E, G) low-LET protons or (F, H) high-LET protons. Clonogenic survival of cells was analysed from two independent experiments (shown by solid and dashed lines). (C, D, G and H) Representative images of colonies in non-irradiated and irradiated plates (the latter of which were seeded with four times the number of cells).

**Fig. 3.**
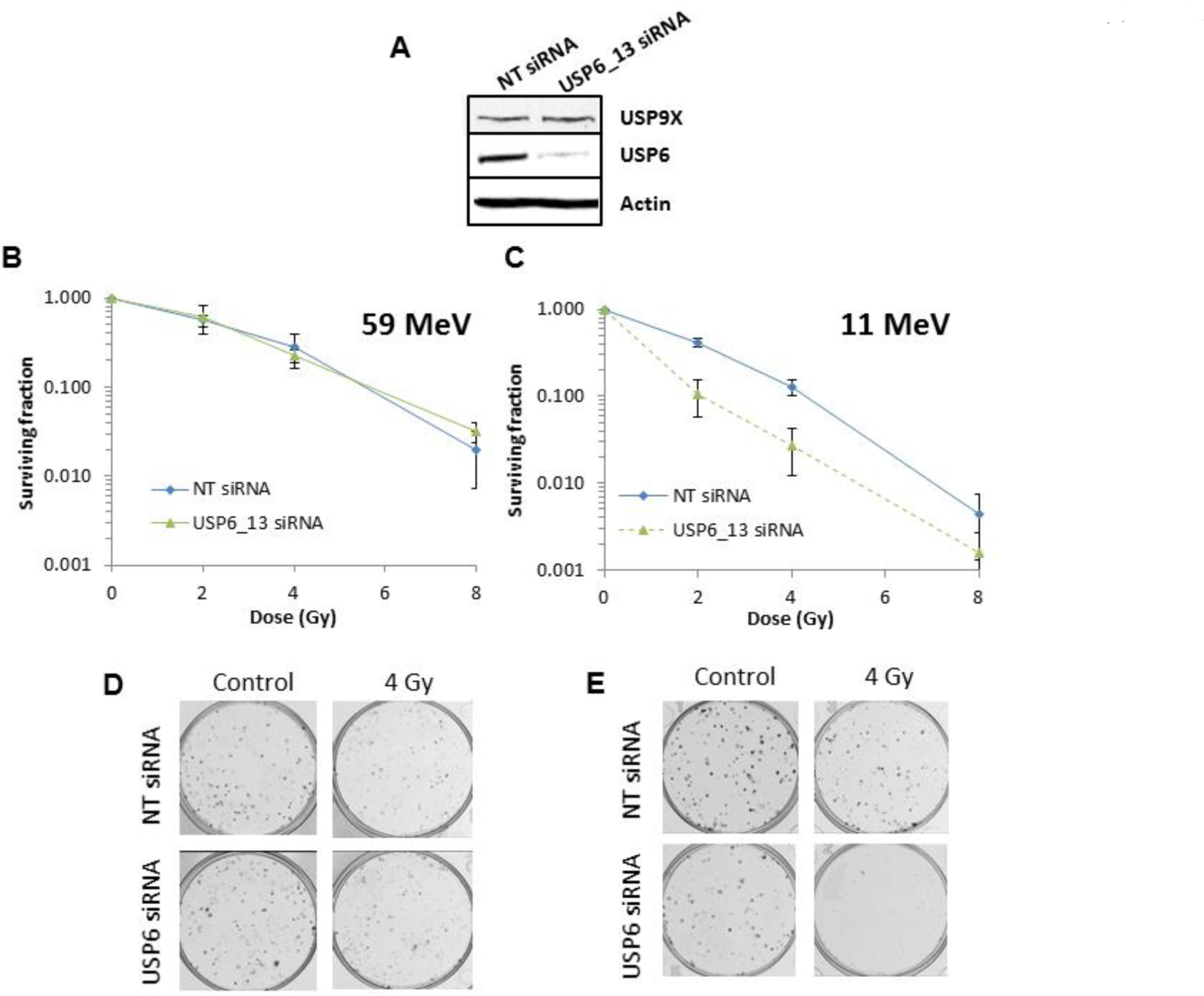
Specific targeting of USP6 leads to enhanced radiosensitivity following high-LET protons. HeLa cells were treated with an individual siRNA sequence targeting USP6 (USP6_13) or a non-targeting (NT) control for 48 h. (A) Whole cell extracts were analysed by immunoblotting. Cells were irradiated with increasing doses of (B, D) low-LET protons or (C, E) high-LET protons. Clonogenic survival of cells was analysed from three independent experiments. Shown is the surviving fraction±S.E. (D and E) Representative images of colonies in non-irradiated and irradiated plates (the latter of which were seeded with four times the number of cells).

### USP6 controls CDD repair and cell cycle progression following high-LET protons

Using an enzyme modified neutral comet assay we found that depletion of USP6 significantly reduces efficiency of CDD repair (**Fig. 4A**, compare blue and yellow bars, and **Fig. 4B**), but not DSB repair, in comparison to NT control siRNA treated cells (**Fig. 4A**, compare green and red bars, and **Fig. 4C**). Also no significant differences in γH2AX and 53BP1 foci, as surrogate markers of DSBs, were observed in cells depleted of USP6 (**Fig. 4D**, **4E**). Cell cycle progression analysis revealed an accumulation of G2/M cells in the absence of irradiation in USP6 siRNA treated cells, which was significantly different from NT siRNA control treated cells at 8-24 h post-irradiation with high-LET protons (**Fig. 5A-C**). Similar cell cycle profiles were observed following low-LET protons, although G2/M arrest in USP6 depleted cells was only significant at 24 h post-irradiation (**Fig. 5D-F**).

**Fig. 4.**
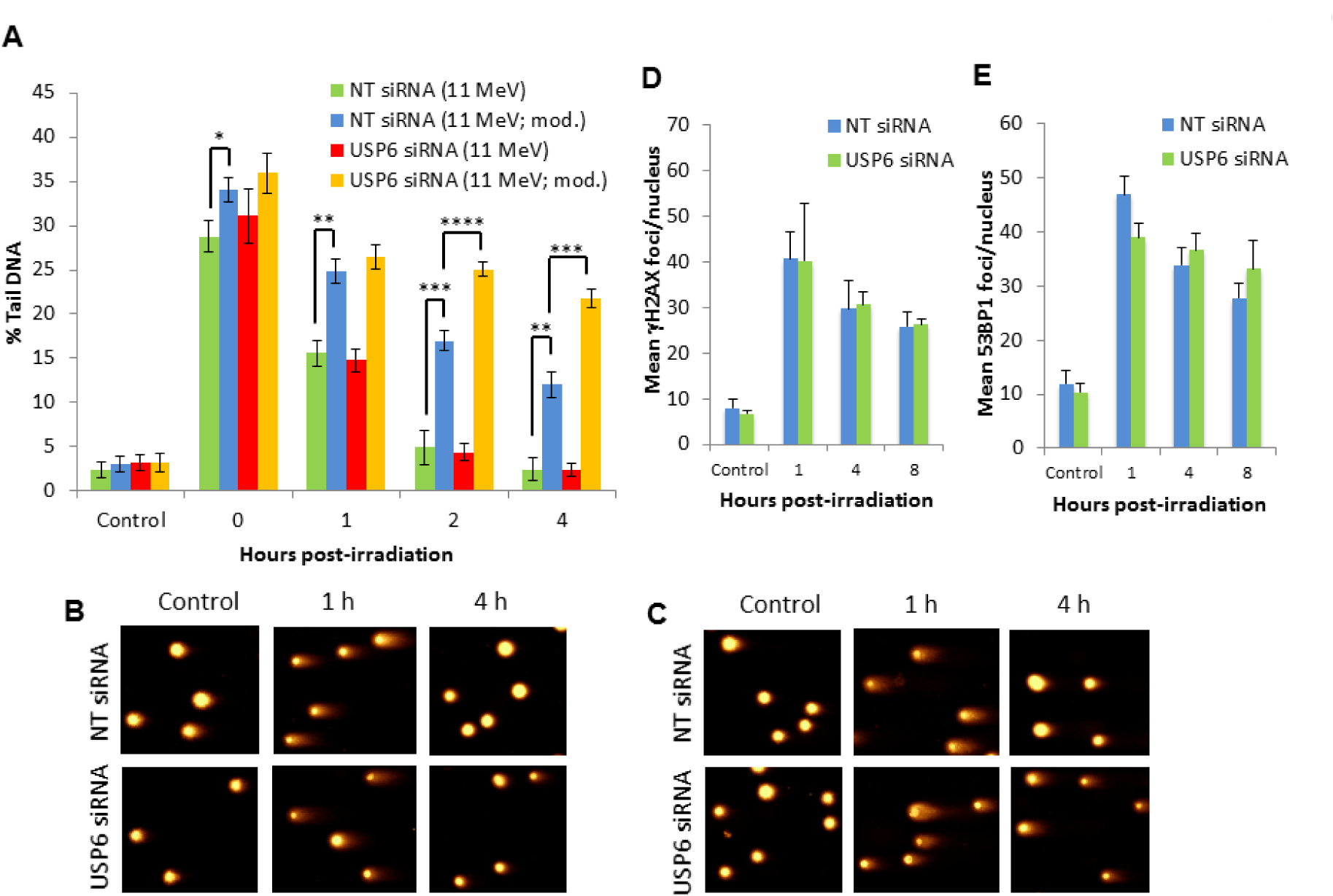
USP6 is required for efficient repair of CDD induced by high-LET protons. HeLa cells were treated with USP6 (USP6_13) siRNA or non-targeting (NT) control siRNA for 48 h. (A-C) Cells were irradiated with 4 Gy high-LET protons and DNA damage measured at various time points post-IR by the enzyme modified neutral comet assay following incubation in the absence (revealing DSBs) or presence (revealing CDD; as indicated by mod) of the recombinant enzymes APE1, NTH1 and OGG1. Shown is the mean % tail DNA±S.D. *p<0.01, **p<0.005, ***p<0.002, ***p<0.001 as analysed by a one sample *t*-test. Representative images of stained DNA in non-irradiated and irradiated cells 1 and 4 h post-irradiation in the presence (B) or absence (C) of recombinant repair enzymes. Cells were irradiated with 4 Gy high-LET protons and (D) γH2AX or (E) 53BP1 foci analysed by immunofluorescent staining at various time points post-IR. Shown is the mean number of foci/nucleus±S.D.

**Fig. 5.**
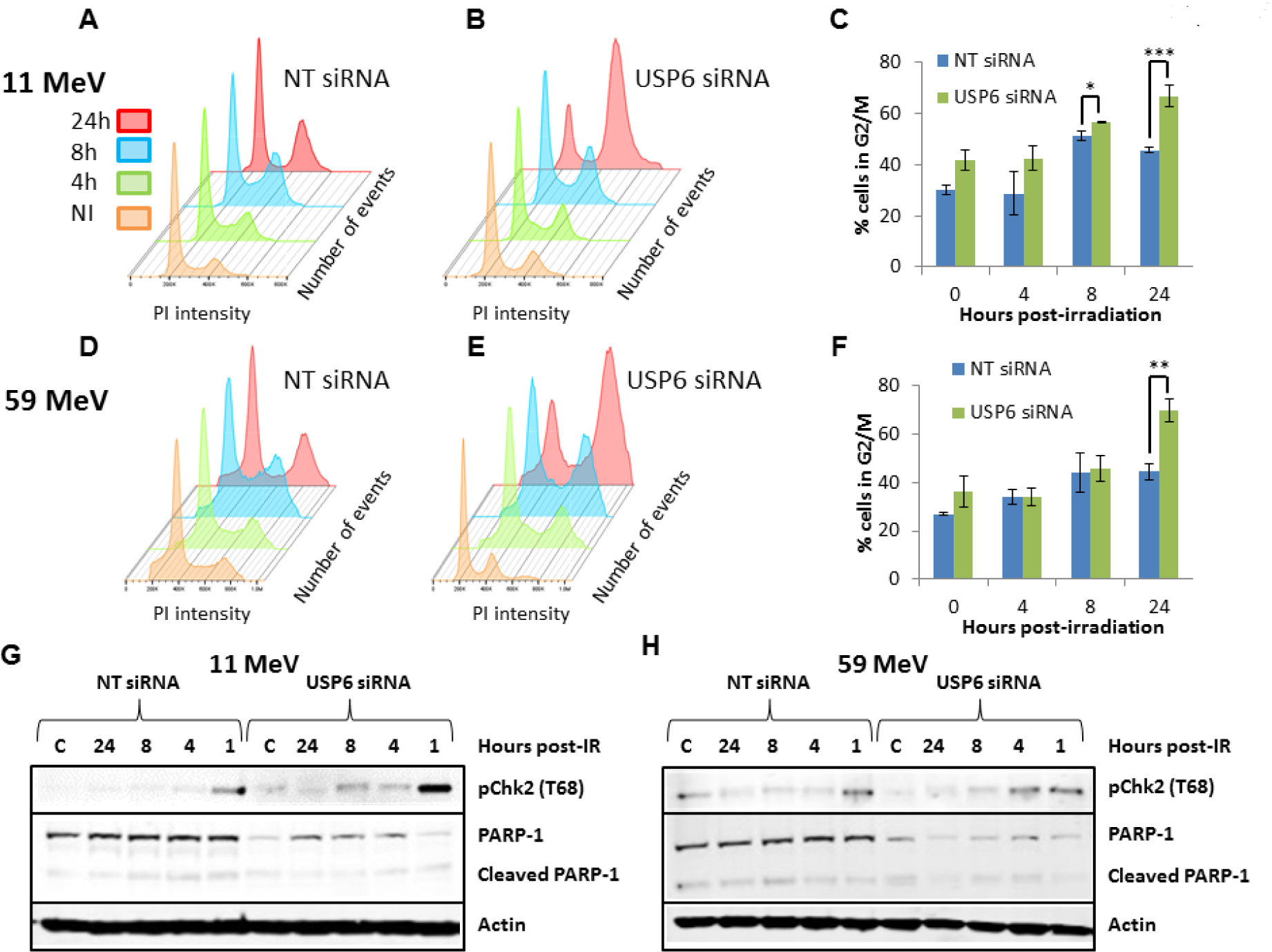
USP6 controls cell cycle progression in response to proton irradiation. HeLa cells were treated with USP6 (USP6_13) siRNA or non-targeting (NT) control siRNA for 48 h. At various time points post-irradiation with 4 Gy (A-C) high-LET or (D-F) low-LET protons, cell cycle profiles were determined by FACS analysis. (C and F) Mean % cells in G2/M phase±S.E. *p<0.05, **p<0.02, ***p<0.01 as analysed by a one sample *t*-test. Cells were collected at various time points post-irradiation with (G) high-LET or (H) low-LET protons and whole cell extracts analysed by immunoblotting.

In USP6 depleted cells, there was a marked increase in Chk2 phosphorylation/activation at 1 h post-irradiation versus NT control siRNA treated cells, which persisted at least up to 8 h post-irradiation consistent with G2/M checkpoint activation (**Fig. 5G**). There was less marked Chk2 phosphorylation/activation in USP6 depleted cells after 1 h post-irradiation with low-LET protons (**Fig. 5H**), suggesting less prominent activation of the G2/M checkpoint. No evidence of significant decreased PARP-1 protein levels, and thus accumulation of cleaved PARP-1 protein as an indicator of apoptosis, in NT control or USP6 siRNA treated cells was evident following either high‐ or low-LET protons (**Fig. 5G, 5H**). However levels of PARP-1 were noticeably lower in USP6 depleted cells.

### USP6 controls PARP-1 protein levels required for resistance to high-LET protons

We substantiated significantly reduced PARP-1 protein levels (~70 %) in USP6 siRNA knockdown cells, in comparison to NT control siRNA cells (**Fig. 6A**). No decreases in XRCC1 and APE1 proteins, also involved in base excision/single strand break repair, were observed in USP6 depleted cells. Cycloheximide-induced inhibition of protein synthesis revealed that PARP-1 is significantly more stable in NT control siRNA cells than in the absence of USP6 (**Fig. 6B, 6C**). To correlate reduced levels, and therefore activity, of PARP-1 in USP6 depleted cells with increased sensitivity to high-LET protons, we treated cells with the PARP inhibitor olaparib. This led to a significant decrease in cell survival in response to high-LET protons versus DMSO treated cells (**Fig. 6D, 6E**) but had no impact on cell survival following low-LET protons (**Fig. 6F, 6G**). Furthermore, olaparib caused a significant delay in repair of CDD-induced by high-LET protons that persisted for at least 4 h post-irradiation versus DMSO treated cells (**Fig. 6H**, compare blue and yellow bars, **Fig. E4A**), phenocopying the effect of USP6 depletion. PARP inhibition had no impact on the global repair of DSBs in response to high-LET protons (**Fig. 6H**, compare green and red bars, **Fig. E4B**).

**Fig. 6.**
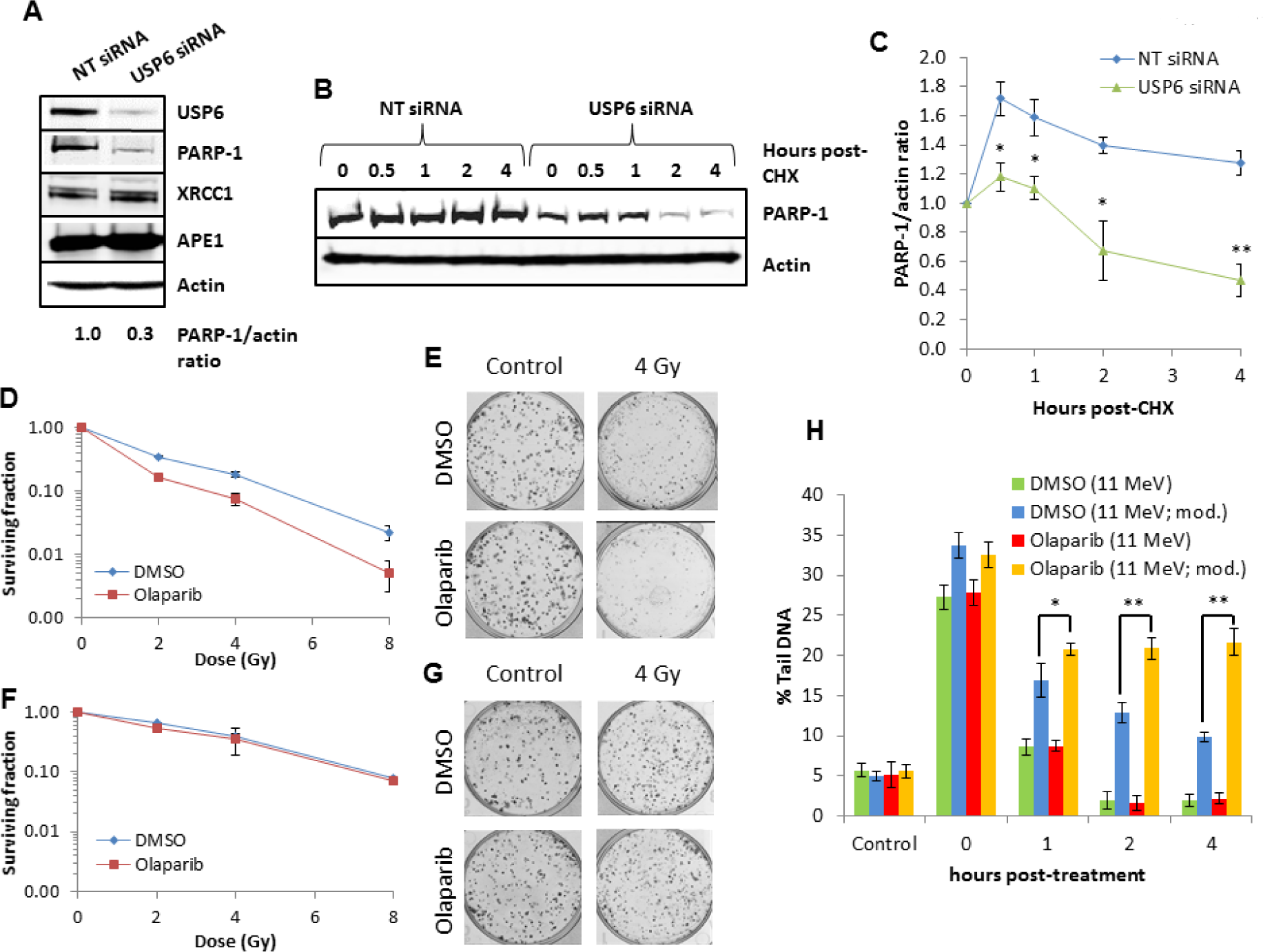
USP6 controls PARP-1 protein levels required for controlling sensitivity to high-LET protons. (A-C) HeLa cells were treated with USP6 (USP6_13) siRNA or non-targeting (NT) control siRNA for 48 h. (A) Whole cell extracts were analysed by immunoblotting. (B and C) Cells were treated with cycloheximide (50 µg/ml) for the time points indicated and whole cell extracts analysed by immunoblotting. (C) Mean PARP-1/actin ratio±S.D. *p<0.005, **p<0.001 as analysed by a two sample *t*-test comparing the PARP-1/actin ratio following USP6 siRNA versus NT control siRNA. (D-G) HeLa cells were treated with DMSO or 1 µM olaparib for 16 h, and irradiated with increasing doses of (D-E) high-LET or (F-G) low-LET protons and clonogenic survival of cells analysed. Shown is the surviving fraction±S.E. (E and G) Respective images of colonies in control and irradiated plates (the latter of which were seeded with four times the number of cells). (H) Cells treated with DMSO or olaparib were irradiated with 4 Gy high-LET protons and DNA damage measured at various time points post-IR by the enzyme modified neutral comet assay following incubation in the absence (revealing DSBs) or presence (revealing CDD; as indicated by mod) of the recombinant enzymes APE1, NTH1 and OGG1. Shown is the mean % tail DNA±S.D. *p<0.01, **p<0.001 as analysed by a one sample *t*-test.

## Discussion

High-LET IR is more effective than low-LET IR at generating CDD which, due to the presence of multiple DNA lesions, represents a challenge to the DNA repair machinery and is a major contributor to IR-induced cell killing. We have recently demonstrated that low energy (relatively high-LET) protons generate increased quantities of CDD in comparison to high energy (low-LET) protons (9). We also highlighted an important role for ubiquitylation in signalling and repair of CDD, mediated by the E3 ubiquitin ligases RNF20/40 and MSL2 that promote histone H2B ubiquitylation which controls cell survival post-irradiation with high-LET protons. We have now utilised an siRNA screen for DUBs to identify and characterise specific enzymes controlling cell survival following high-LET IR. We demonstrated that depletion of USP6 causes significantly increased radiosensitivity to high-LET IR (low energy protons and α-particles) but not low-LET IR (high energy protons or x-ray irradiation), which is mediated by instability of PARP-1 protein levels, a subsequent deficiency in CDD repair, and G2/M cell cycle checkpoint arrest. The importance of PARP-1 was corroborated by using olaparib, which mimics the effects of USP6 depletion.

Previous studies have utilised both siRNA screening and protein overexpression to examine the roles of DUBs in the cellular response to IR-induced DNA damage, particularly DSBs (15-17). These are in contrast to our study which focussed on DUBs responsive to high-LET IR that generates increasing amounts of CDD, and the comparison to low-LET IR, but also we utilised cell survival as an end-point. Moreover, we have specifically examined the impact of proton irradiation on cell survival. Interestingly our siRNA screen revealed that depletion of a greater number of DUBs increased cellular radiosensitivity following low-LET protons in comparison to high-LET protons (30 and 2 DUBs, respectively yielded a further >50 % reduction in cell survival), suggesting that the cell killing effects of high-LET protons are difficult to exacerbate. In contrast, enhancing the radiosensitivity of cells following low-LET protons would appear much simpler to achieve via depletion of a number of DUBs. Another important observation is the differences in specific DUB enzymes whose depletion leads to altered radiosensitivity in response to low-LET protons versus x-ray irradiation, although we have to be cautious in drawing conclusions from screens involving just a single experiment. Focussing on DUBs whose depletion led to selective radiosensitisation of cells following high-LET versus low-LET protons, we validated that USP6 plays a major role in this process. Previous reports on the cellular role and targets of USP6 are sparse, although it has been suggested to deubiquitylate Frizzled and promote Wnt signalling (18), and to deubiquitylate Jak1 causing activation of the STAT3 signalling pathway (19). USP6 has also been demonstrated to regulate the stability of the c-Jun transcription factor (20), and high USP6 protein expression has been observed in bone and soft tissue tumours (21). We now extend these observations by demonstrating that USP6 plays a critical cellular role in controlling radiosensitivity to high-LET IR and in the cellular DNA damage response. Indeed this effect appears to be achieved by reducing the efficiency of CDD repair as a consequence of a significant reduction in PARP-1 protein levels and stability, and a G2/M cell cycle arrest. Importantly we showed that cells treated with the PARP inhibitor olaparib, also displayed increased radiosensitivity to high-LET protons as a consequence of deficiencies in CDD repair. This would suggest synergy between PARP inhibition and CDD-induced by high-LET IR in promoting cancer cell killing, similar to the concept of synthetic lethality observed with PARP inhibitors in combination with BRCA (DSB repair)-deficient cancers (22,23). This data correlates with our previous evidence demonstrating that CDD-induced by high-LET protons appear to be largely SSB in nature and thus are dependent on PARP-1, and predictably other SSB repair proteins, for repair (9). Whilst PARP-1 is known to be regulated by ubiquitylation (11), particularly the poly(ADP-ribosyl)ated form of the protein by the E3 ubiquitin ligases Iduna/RNF146 (24) and CHFR (25), to our knowledge the identification of a specific DUB involved in controlling PARP-1 protein levels has not previously been reported. Therefore our data would suggest an interplay between USP6 and Iduna and/or CHFR in the controlled regulation of PARP-1 protein levels, which requires further clarification. Nevertheless, our evidence now describes that USP6 and PARP-1 should be considered as important factors in the responsiveness of cancer cells to radiotherapy, particularly proton beam therapy that at the distal end of the Bragg peak can generate high-LET protons.

## Acknowledgement

–The authors thank Prof T. Carey for providing the UMSCC cells and Prof G. Dianov for providing APE1, NTH1 and OGG1 bacterial expression plasmids, and XRCC1 and APE1 antibodies. We also thank Brian Marsland, Andy Wray and Ian Taylor at the Clatterbridge Cancer Centre for technical assistance with proton irradiation of cells.

